# Physiological Growth Indices of Maize (*Zea mays* L.) Genotypes in Sylhet

**DOI:** 10.1101/518993

**Authors:** M. Tazul Islam, A.F.M. Saiful Islam, M. Sharaf Uddin

**Author notes:** Corresponding author and Professor, Department of Agro forestry and Environmental Science, Sylhet Agricultural University, Sylhet-3100, Bangladesh.

## Abstract

Maize (*Zea mays* L.) is an important food and feed crop in Bangladesh as well as in the world. But its cultivation is limited in the soils of Sylhet, Bangladesh. To study the physiological growth indices of maize genotypes an experiment was carried out at the experimental field of Sylhet Agricultural University (SAU), Bangladesh during December 2014 to May 2015. The maize genotypes as the experimental materials were ZM 0001, ZM 0002, ZM 0003, ZM 0004, ZM 0005, ZM 0006 ZM 0007 and BARI Maize 6. The experiment was laid out in a Randomized Complete Block Design (RCBD) with three replications. Seeds were sown on 26 December 2014. Proper cultural management practices were followed as and when necessary during the growing period of the crop. Data were recorded on dry matter partitioning, different growth rate parameters and dry matter production of different plant parts at 30 days interval. The results indicated significant variations in almost all of the growth indices. Dry matter accumulation continued to increase from seedling to harvesting. The highest root dry matter was found in ZM 0007 (36.03 g) whereas the highest shoot and leaf dry matter was found in BARI Maize 6 (254.6 and 45.99 g, respectively) at harvest. The highest total dry matter (TDM) was produced in BARI Maize 6 at harvest (287.5 g) and AGR and CGR value was also highest in BARI Maize 6 (5.686 g day^-1^ and 45.48 g m^-2^ day^-1^) but at 60-90 days after germination (DAG). The highest RGR value was in ZM 0001 at seedling stage (114.1 mg g^-1^ day^-1^). A zigzag pattern graph was observed in all genotypes for RSR, RWR and SWR at different growth stages but a declining trend from seedling to harvesting was observed for LWR. Correlation between the growth rate parameters and kernel yield showed that shoot, leaf, total dry weight, AGR, CGR, RGR and SWR had significant positive correlation with yield.

## Introduction

Maize *(Zea mays* L.) is an important cereal crop of the world. It is ranked as the third most important cereal crops followed by rice and wheat in the world (FAO, 2010). It is used as staple food in many parts of the world. As a cereal crop it can easily be grown successfully under rainfed condition, requires little management practices and less amount of capital which can meet the additional food requirement of the world. Moreover, being a C4 crop plant, having a large leaf area it is more efficient in converting solar energy to dry matter than most other cereals which can give high biological yield as well as grain yield relatively in a shorter period of time due to its unique photosynthetic mechanism (Hatch and Slack, 1966). Beside this, it is a multipurpose crop which provides food for human beings, feed and fodder for poultry and livestock. In terms of nutrition, it has high nutritional value as it contains about 72% starch, 10% protein, 4.8% oil, 8.5% fibre, 3.0% sugar and 1.7% ash (Chaudhary, 1983). In Bangladesh maize is mainly used as food (Rooti, Chapra, Firni, Satu etc.), feed (poultry, dairy, and fish), Green cob, boiled, roasted and pop, fodder and fuel. But it is mainly used in poultry industry which requires 90% of total annual maize demand (Hasan, 2008). The peoples of Bangladesh are habituated in rice based food system. But the average per unit production of rice is lower than the maize (BBS, 2011). Maize is also better than the rice and wheat in terms of nutritional value (Rohman *et al.*, 2011). So if the traditional rice based food habit can be diversified using maize it can play a vital role in the food security of Bangladesh. At present, the annual requirement for maize in Bangladesh is 1.6 million metric tons (Ahmed, 2013). But only 1.02 million metric ton maize is produced in the country per annum (BBS, 2011). So, Bangladesh needs more maize production to meet its annual demand for various uses which push forward for increased maize cultivation. Physiological growth analysis is a way to assess what events occurs during plant growth and eventually it is important in the prediction of yield of crop (Hokmalipour and Darbandi, 2011). Biomass partitioning is important for crop production and plants usually divert accumulated biomass to other plant parts to ensure and maintain a high production capacity (Srivastava and Gaiser, 2008). It is one of the tactics to analyze the yield-influencing factors and plant development (Berzsenyi and Lap, 2004). So, plant growth analysis is considered as crucial in the development of productive lines and varieties. Therefore, an experiment was conducted to evaluate the variability of biomass accumulation, plant growth rate at different growth stages and to correlate different growth rate parameters with the yield of eight maize genotypes.

## Materials and Methods

The experiment was conducted at the experimental field of Department of Crop Botany and Tea production Technology, Sylhet Agricultural University (SAU), Bangladesh from December 2014 to May 2015. Eight maize genotypes viz. ZM 0001, ZM 0002, ZM 0003, ZM 0004, ZM 0005, ZM 0006, ZM 0007 and BARI Maize-6 (control) were used as experimental materials. Among them, BARI Maize-6 was collected from Bangladesh Agricultural Research Institute (BARI) and the rests of the genotypes were collected from abroad. The experiment was laid out in a randomized complete block design (RCBD) with three replications. The unit plot size was 2.5 m × 1.5 m, row to row and plant to plant distance were 50 cm and 25 cm, respectively. Crop husbandry was maintained according to BARI recommendation to ensure optimum plant growth and development. In order to determine dry matter, five plants were randomly selected from each plot and uprooted for collecting necessary data. The plant parts were separated into root, shoot and leaves. Then the samples were kept into an oven at 80 ± 2 °C for 72 hours and the respective dry weights were recorded. The procedure was followed every 30 days interval from seed germination to the final harvest. Absolute growth rate (AGR), Crop growth rate (CGR), Relative growth rate (RGR) were determined using the following formulas (Aliabadi *et al,* 2008):

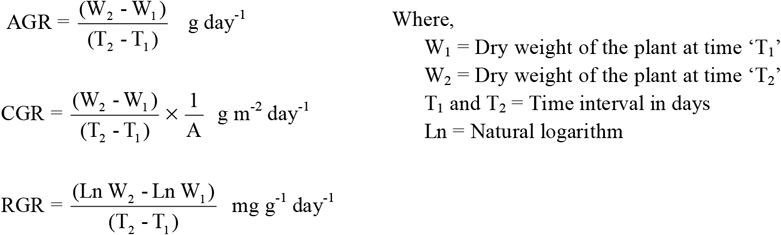

Root shoot ratio (RSR), Root weight ratio (RWR), Shoot weight ratio (RWR) and Leaf weight ratio (RWR) were calculated using the following formula:

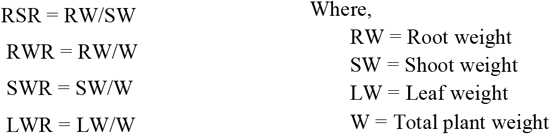

## Results and Discussion

### Root dry weight (g)

The dry weight of roots of different genotypes showed significant variation at 30 and 60 DAG and didn’t show any variation at 90 DAG and at harvest [Fig. 1(a)]. The highest root dry weight was produced in ZM 0007 (36.03 g) at harvest whereas the lowest was produced in ZM 0006 (31.83 g). It is observed that the dry weight of roots increased gradually from 30 to 60 DAG and 90 DAG to harvesting. But from 60 to 90 DAG it increased rapidly as compared to other growth stages. This was due to rapid increase in number and length of roots at this stage.

### Shoot dry weight (g)

The data presented on shoot dry weight showed significant variations at all growth stages [Figure 1(b)]. The highest shoot dry weight was produced in BARI Maize 6 (254.6 g) at harvest which was statistically similar to ZM 0005 (244.7 g) and ZM 0004 (239.9 g). On the other hand, the minimum shoot dry weight was produced in ZM 0001 (216.8 g) which was statistically identical to ZM 0002 (219.7 g). Shoot dry weight increased gradually at 30 to 60 DAG and 90 DAG to harvesting. But from 60 to 90 DAG it was increased sharply as compared to other growth stages. This was due to appearance of tassel and ears in the plants. Beside this, at this stage the height, stem girth and number of leaves of the plant increased rapidly. The variation among the genotypes in terms of shoot dry weight was due to the variation in size of tassel and ears as well as variation in plant height, stem girth and number of leaves of the genotypes.

### Leaf dry weight (g)

The dry weight of leaves of different genotypes revealed significant variations at all growth stages [Figure 1(c)]. The highest leaf dry weight was produced in the genotype BARI Maize 6 (45.99 g) at harvest which was statistically identical to ZM 0005 (43.45 g), ZM 0003 (42.99 g) and ZM 0007 (42.55 g), while minimum leaf dry weight was produced in ZM 0001 (33.84 g). It is observed that the dry weight of leaves was increased at a constant rate from 30 to 90 DAG. It was due to the increase in number, length and breadth of leaves. After 90 DAG, the rate of increase reduced drastically even there was reduction in leaf weight in the genotypes ZM 0001, ZM 0003 and ZM 0004. The reason behind this was abscission of leaves at maturity stage. The variation among the genotypes for leaf dry weight was due to the difference among the genotypes for number and size of leaves.

### Total dry matter (g)

In respect of total dry matter, there was a significant difference amongst the genotypes [Fig. 1 (d)]. BARI Maize 6 was superior for dry matter production as it produced the highest amount of dry matter (287.5 g) at harvest which was statistically similar to ZM 0005 (279 g), while the lowest dry matter was produced in ZM 0001 (250.1 g). Dry matter production rate was increased till harvesting. But the rate of increment at 60 to 90 DAG was faster as compared to other growth stages. This was due to rapid increase in plant height, stem girth, root number, length and as well as increase in leaf number and size. Beside this, the appearance of tassel and ear at this stage contributed greatly to the sharp increment of total dry matter. The variation amongst the genotypes for total dry matter was due to the differences among the genotypes for weight of root, shoot and leaves. This result is in agreement with the findings of Ibeawuchi *et al* (2008) who reported that the highest plant biomass was produced at taselling and silking stage of maize cultivars.

**Fig. 1.**
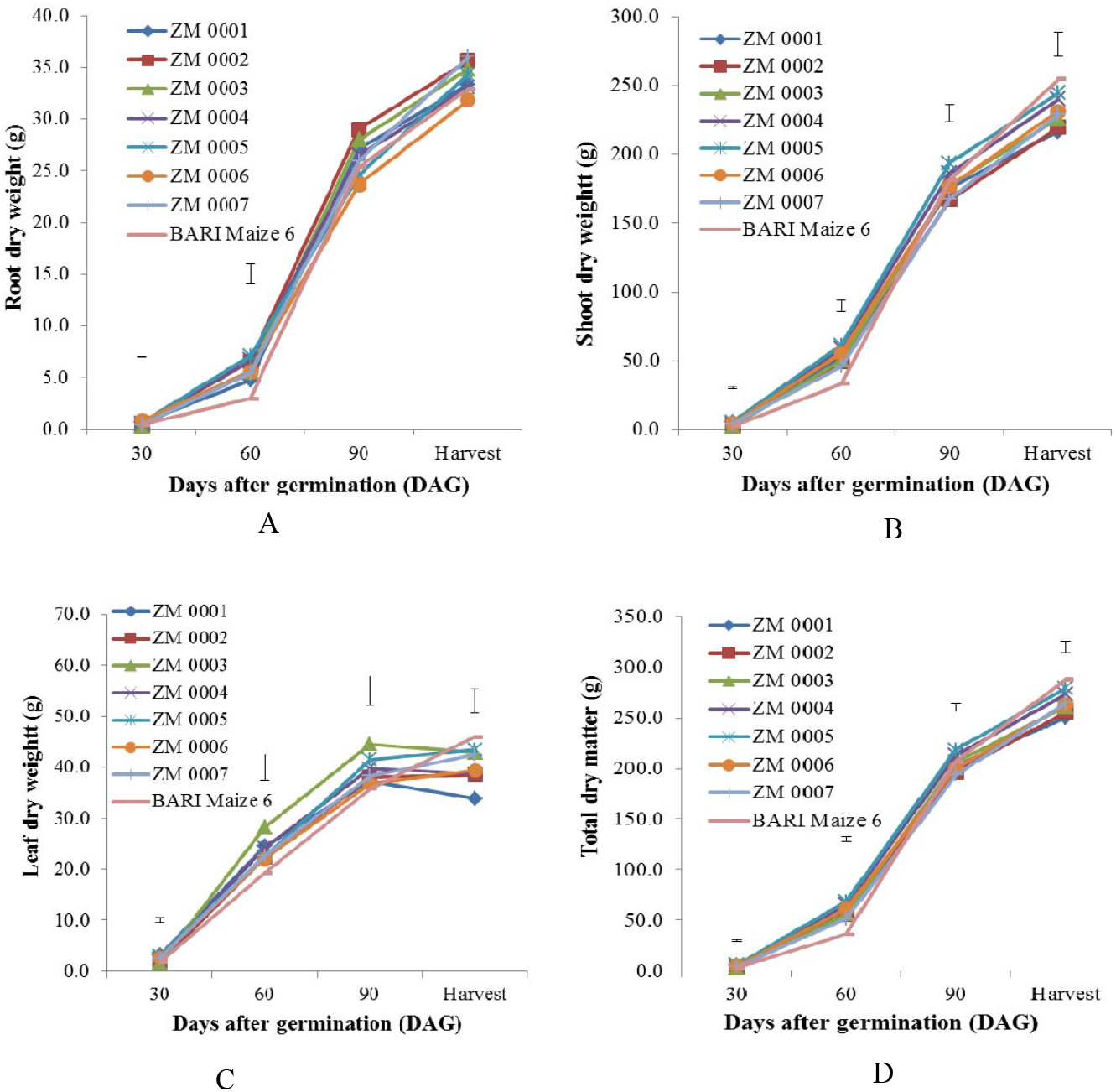
Weight of A) Root, B) Shoot, C) Leaf and D) Total dry matter of eight maize genotypes at different days after germination [Vertical bars represent lsd_(0.05)_]

### Crop growth rate (CGR)

There were significant differences among the genotypes in terms of crop growth rate (CGR) [Fig. 2(b)]. The highest growth rate was obtained in BARI Maize 6 (45.48 g m^-2^ day^-1^) at 60-90 DAG and minimum was in ZM 0006 (37.17 g m^-2^ day^-1^) which was closely followed by rest of the genotypes. The rate of the crop growth was increased up to 90 DAG and then it was decreased sharply till harvest. The reason behind the increment of crop growth rate was due to accumulation of more dry matter by the plants. After 90 DAG it declined because after vegetative stage dry matter accumulation increases but the rate of accumulation was not as much as vegetative stage. The variation among the genotypes was also due to the variation in dry matter accumulation by different genotypes. The similar results were also found by Hokmalipour and Darbandi (2011), Aliu (2010) and Limpinuntana *et al.* (2010). Limpinuntana *et al.* (2010) reported that during the early period especially after 30 DAG, the CGR increases sharply until 90 DAG then it gradually decreases.

**Fig. 2.**
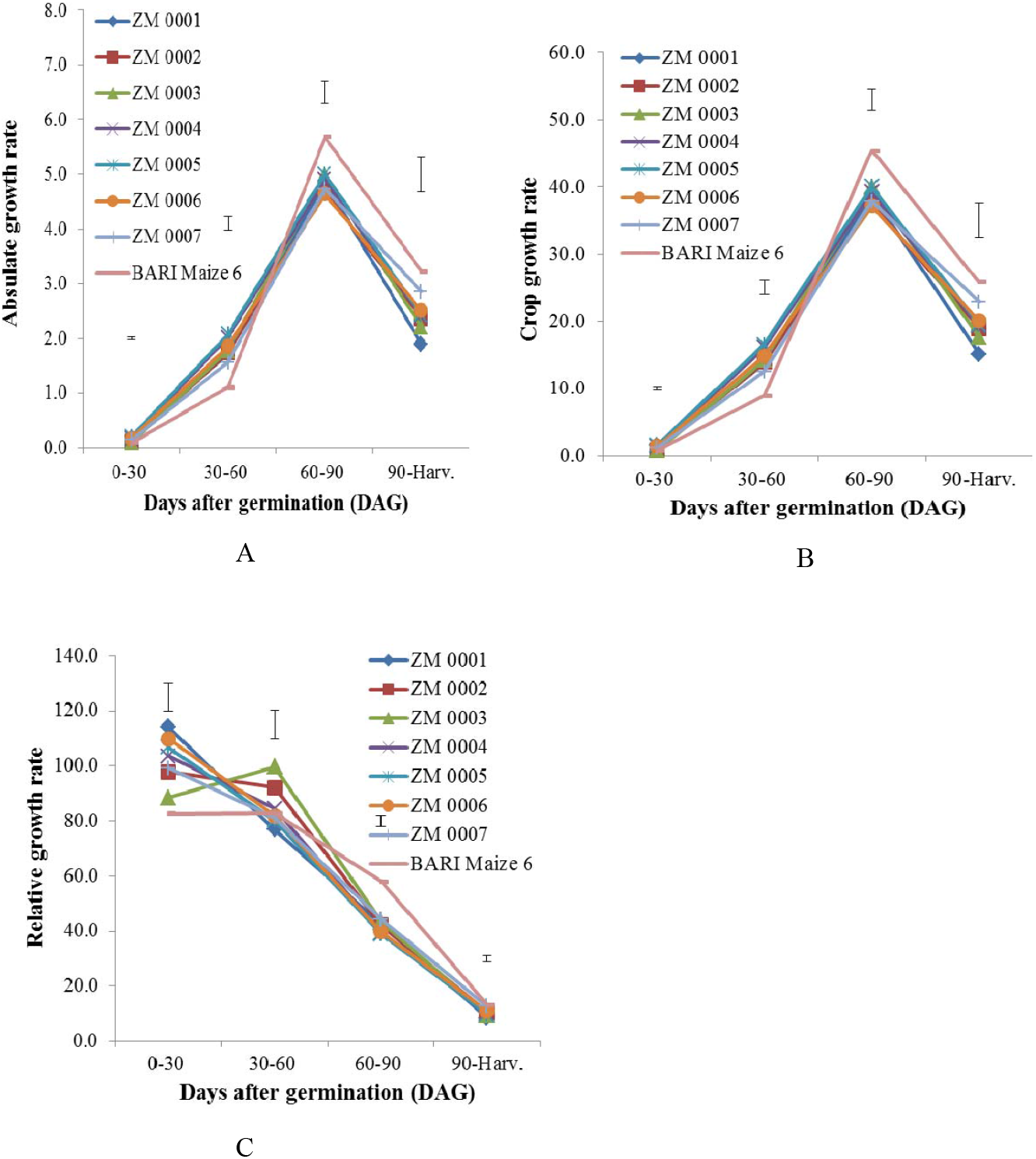
A) Absolute growth rate (AGR), B) Crop growth rate (CGR) and C) Relative growth rate (RGR) of eight maize genotypes at different days after germination [Vertical bars represent lsd(0.05)]

### Relative growth rate (RGR)

Relative growth rate (RGR) of different genotypes exposed significant variations at different growth stages [Fig. 2(c)]. The highest value of RGR was found in ZM 0001 (114.1 mg g^-1^ day^-1^) which was closely followed by ZM 0006 (109.9 mg g^-1^ day^-1^), ZM 0005 (106.7 mg g^-1^ day^-1^) and ZM 0004 (103.6 mg g^-1^ day^-1^). On the other hand, the lowest value was found in BARI Maize 6 (82.57 mg g^-1^ day^-1^). It was observed that the RGR value was the highest at early growth stage and then it tends to decrease until harvest. The variation among the genotypes was due to the variation in dry matter accumulation by different genotypes. These results are in conformity with the findings of Hokmalipour and Darbandi (2011) who reported that relative growth rate (RGR) was significantly different among the maize cultivars and observed a declining trend of RGR as the crop proceeds towards maturity.

### Root shoot ratio (RSR)

The data on root shoot ratio (RSR) presented in Figure 3(a) indicated significant variations among the genotypes at all growth stages except at 60 DAG. The highest value of root shoot ratio was found in ZM 0006 (0.1818) at 30 DAG which was statistically similar to BARI Maize 6 (0.1739) and ZM 0002 (0.1629) and minimum ratio was found in ZM 0007 (0.1051) which was closely followed by ZM 0004 (0.109) and ZM 0005 (0.1105). It was observed that the graph of RSR value of all genotypes showed a zigzag pattern. This was due to the variation in dry weight of roots and shoots. The variation within the genotypes was also due to the variation in dry weight of roots and shoots.

### Root weight ratio (RWR)

The genotypes revealed significant variations among them at all growth stages except at 60 DAG for root weight ratio (RWR) [Fig. 3(b)]. The highest ratio was recorded in the genotype ZM 0006 (0.1536) at 30 DAG which was statistically similar to BARI Maize 6 (0.1481) and ZM 2 (0. 14) and minimum ratio was found in ZM 0007 (0.0951) which was closely followed by ZM 0004 (0.0982) and ZM 0005 (0.09933). It was observed that the graph of RWR value of all genotypes showed a zigzag pattern. This was due to the variation in dry weight of roots and total dry matter. The variation within the genotypes was also due to the variation in dry weight of roots and total dry matter.

**Fig. 3.**
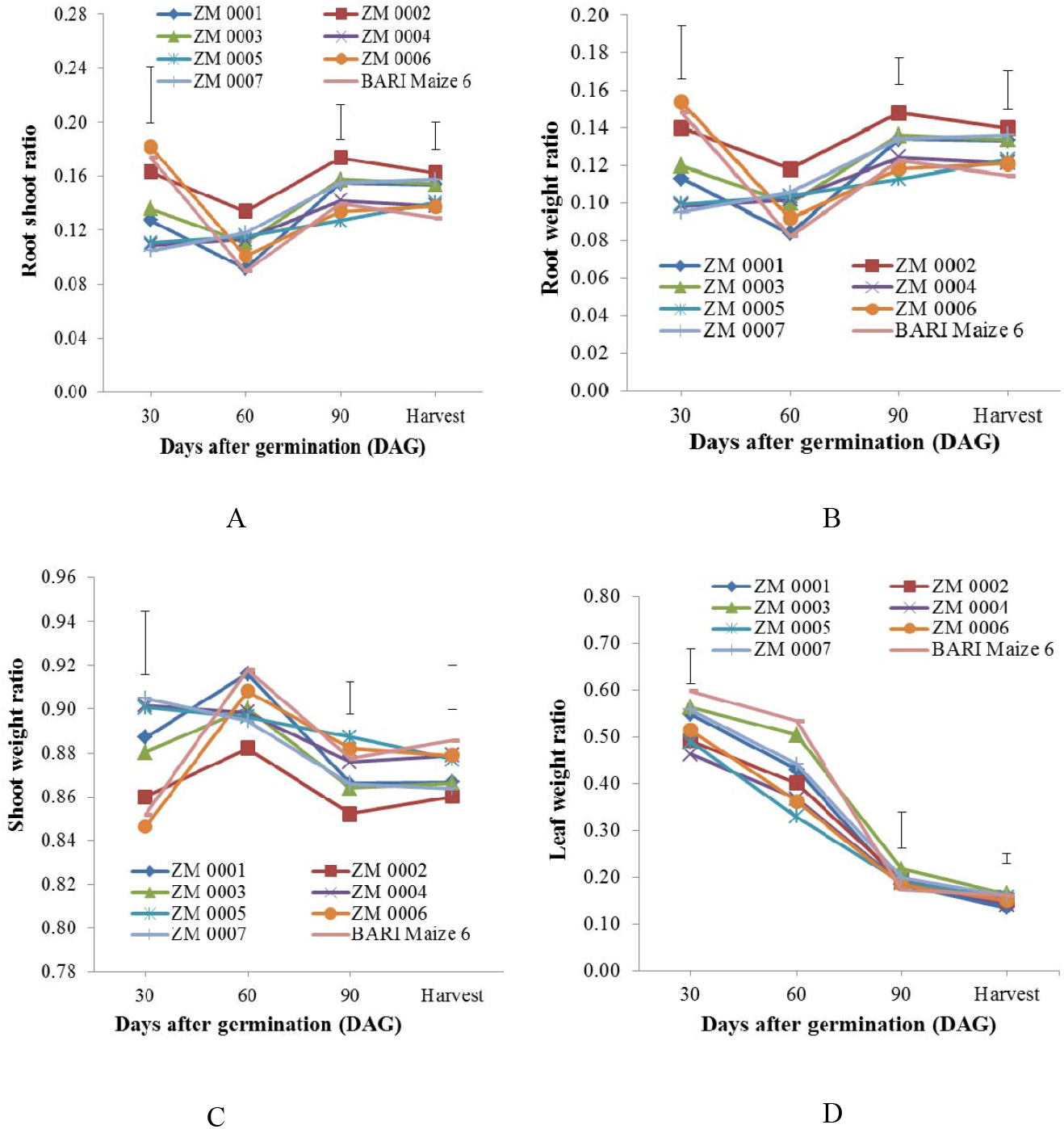
A) Root shoot ratio (RSR), B) Root weight ratio (RWR), C) Shoot weight ratio (SWR) and D) Leaf weight ratio (LWR) of eight maize genotypes at different days after germination [Vertical bars represent lsd_(0.05)_]

### Shoot weight ratio (SWR)

There were significant variations among the maize genotypes for shoot weight ratio (SWR) at all growth stages except at 60 DAG [Fig. 3(c)]. The highest value for SWR was found in BARI Maize 6 (0.918) while minimum shoot weight ratio was in ZM 0002 (0.882). It was observed that the graph of SWR value of all genotypes showed a zigzag pattern. This was due to the variation in dry weight of shoots and total dry matter. The variation within the genotypes was also due to the variation in dry weight of shoots and total dry matter.

### Leaf weight ratio (LWR)

Leaf weight ratio (LWR) of eight maize genotypes showed significant variations among them at all growth stages except at 60 DAG [Fig. 3(d)]. The highest value for LWR was found in BARI Maize 6 (0.5967) which was statistically identical to ZM 0003 (0.5635), ZM 0007 (0.5579) and ZM 0001 (0.548) while minimum leaf weight ratio was in ZM 0004 (0.4624). It was observed that the leaf weight ratio tends to decrease from early growth stage towards harvest. This was due to increase of total dry matter compared to leaf dry matter. The variation among the genotypes was also due to the variation in dry weight of leaves and total dry matter.

### Correlation between growth characters (harvesting stage) and grain yield of different maize genotypes

Correlation between growth characters and grain yield presented in Table 1 represents that grain yield was significantly positively correlated with shoot dry weight (0.84), leaf dry weight (0.62), total dry weight (0.82), AGR (0.70), CGR (0.70), RGR (0.55) and SWR (0.62). It meant that grain yield would increase with the increase of these characters. Root shoot ratio (RSR) (−0.62) and RWR (−0.62) showed significant negative correlation with grain yield. It meant that with the increase of these characters grain yield would decrease. Leaf weight ratio (LWR) had positive but non-significant correlation with grain yield (0.25) and root dry weight had negative non-significant correlation with grain yield (−0.07).

**Table 1.** Correlation matrix between growth characters (harvesting stage) and grain yield of different maize genotypes

| Characters | Shoot dry weight | Leaf dry weight | Total dry weight | AGR | CGR | RGR | RSR | RWR | SWR | LWR | Grain yield |
| --- | --- | --- | --- | --- | --- | --- | --- | --- | --- | --- | --- |
| Root dry weight | 0.01<sup>NS</sup> | -0.01<sup>NS</sup> | 0.17<sup>NS</sup> | 0.22<sup>NS</sup> | 0.22<sup>NS</sup> | 0.22<sup>NS</sup> | 0.73** | 0.73** | -0.73** | -0.10<sup>NS</sup> | -0.07<sup>NS</sup> |
| Shoot dry weight |  | 0.57** | 0.99** | 0.70** | 0.70** | 0.50* | -0.68** | -0.67** | 0.67** | 0.11<sup>NS</sup> | 0.84** |
| Leaf dry weight |  |  | 0.56** | 0.55** | 0.55** | 0.48* | -0.40<sup>NS</sup> | -0.40<sup>NS</sup> | 0.40<sup>NS</sup> | 0.88** | 0.62** |
| Total dry weight |  |  |  | 0.73** | 0.73** | 0.53** | -0.55** | -0.55** | 0.55** | 0.10<sup>NS</sup> | 0.82** |
| AGR |  |  |  |  | 1.00** | 0.96** | -0.31<sup>NS</sup> | -0.31<sup>NS</sup> | 0.31<sup>NS</sup> | 0.24<sup>NS</sup> | 0.70** |
| CGR |  |  |  |  |  | 0.96** | -0.31<sup>NS</sup> | -0.31<sup>NS</sup> | 0.31<sup>NS</sup> | 0.24<sup>NS</sup> | 0.70** |
| RGR |  |  |  |  |  |  | -0.17<sup>NS</sup> | -0.17<sup>NS</sup> | 0.17<sup>NS</sup> | 0.28<sup>NS</sup> | 0.55** |
| RSR |  |  |  |  |  |  |  | 1.00** | -1.00** | -0.15<sup>NS</sup> | -0.62** |
| RWR |  |  |  |  |  |  |  |  | -1.00** | -0.15<sup>NS</sup> | -0.62** |
| SWR |  |  |  |  |  |  |  |  |  | 0.15<sup>NS</sup> | 0.62** |
| LWR |  |  |  |  |  |  |  |  |  |  | 0.27<sup>NS</sup> |

\* $P < 0.05$ , \*\* $P < 0.01$ , \*\*\* $P < 0.001$ : probability from $t$ -test between growth characters at harvesting stage and grain yield.

Root dry weight showed significant positive correlation with RSR (0.73), RWR (0.73) but that of significantly negative with SWR (−0.73). Shoot dry weight had significant positive correlation with leaf dry weight (0.57), total dry weight (0.99), AGR (0.70), CGR (0.70), RGR (0.50) and SWR (0.67) but that of significantly negative with RSR (−0.68) and RWR (−0.67). Leaf dry weight showed significant positive correlation with total dry weight (0.56), AGR (0.55), CGR (0.55), RGR (0.48) and LWR (0.88). Total dry weight revealed significant positive correlation with AGR (0.73), CGR (0.73), RGR (0.53) and SWR (0.55) but that of significantly negative with RSR (−0.55) and RWR (−0.55).

AGR had significant positive correlation with CGR (1.00) and RGR (0.96). CGR showed significant positive correlation with RGR (0.96). RSR had significant positive correlation with RWR (1.00) but that of significantly negative with and SWR (−1.00). RWR had significant negative correlation with SWR (−1.00).

## Conclusion

It was revealed from the study that BARI maize 6 performed best among the studied genotypes in respect of dry matter accumulation and growth rate. But ZM 0004, ZM 0005, ZM 0006 and ZM 0007 also showed good potentials in respect of dry matter accumulation and growth rate parameters. So these genotypes could be used as potential material in the future research works.

